# SWR1 Chromatin Remodeling Complex Prevents Mitotic Slippage during Spindle Position Checkpoint Arrest

**DOI:** 10.1101/749440

**Authors:** Ayse Koca Caydasi, Anton Khmelinskii, Zoulfia Darieva, Bahtiyar Kurtulmus, Michael Knop, Gislene Pereira

## Abstract

Faithful chromosome segregation in budding yeast requires correct positioning of the mitotic spindle along the mother to daughter cell polarity axis. When the anaphase spindle is not correctly positioned, a surveillance mechanism, named as the spindle position checkpoint (SPOC), prevents the progression out of mitosis until correct spindle positioning is achieved. How SPOC works on a molecular level is not well-understood. Here, we performed a genome-wide genetic screen to search for components required for SPOC. We identified the SWR1 chromatin-remodeling complex (SWR1-C) among the several novel factors that are essential for SPOC integrity. Cells lacking SWR1-C were able to activate SPOC upon spindle misorientation but underwent mitotic slippage upon prolonged SPOC arrest. This mitotic slippage required the Cdc14-early anaphase release pathway and other factors including the SAGA histone acetyltransferase complex, proteasome components, the mitotic cyclin-dependent kinase inhibitor Sic1 and the mitogen-activated protein kinase Slt2/Mpk1. Together, our data establish a novel link between chromatin remodeling and robust checkpoint arrest in late anaphase.

**AUTHORS SUMMARY:** Before it physically divides into two, the cell must duplicate its genetic material and separate the duplicated copies to the opposite poles of the cell with the help of the spindle machinery. The direction along which the genetic material is separated has different consequences on cell division, especially when the opposite poles of the cell differ from each other, as is the case of asymmetric cell division. Every cell division in budding yeast is asymmetric. The new (daughter) cell grows on the old (mother) cell and pinches of from this location at the end of the cell division, giving rise to a new and an old cell. The daughter and mother cells differ in size and composition, thus the cell division is asymmetric. In order for the daughter cell to receive one copy of the duplicated genetic material, budding yeast has to separate the copies of its genetic material along the mother to daughter cell direction, which is possible by placing the spindle apparatus along this direction.

A surveillance mechanism named the Spindle Position Checkpoint (SPOC) in budding yeast monitors the position of the mitotic spindle and prevents cells from dividing if the spindle fails to align in the mother to daughter direction. The cell can resume cell division only after correcting the position of the spindle followed by inactivation of SPOC. This way SPOC prevents multi-nucleation and enucleation, and hence it is a crucial mechanism to maintain the correct ploidy. It has been known that about five proteins play a role in positively supporting the SPOC. Yet, how SPOC works on a molecular level remains ill understood.

In this study, we aimed to find out novel components of SPOC. Through an unbiased genome-wide genetic screen, we successfully identified several new components of the SPOC machinery. Among several other novel proteins identified, we investigated the role of the SWR1 chromatin remodeling complex (SWR1-C) in more detail. We show that the SWR1-C has a function in preventing cells with mis-positioned spindles from resuming cell division when the spindle stays mis-positioned for a prolonged time (mitotic slippage). Our data indicated that SWR1-C is not required to start the immediate SPOC response, rather it is important to keep the prolonged SPOC arrest.

## INTRODUCTION

In budding yeast *Saccharomyces cerevisiae*, the mitotic spindle must align along the mother-daughter cell polarity axis to deliver one set of chromosomes to the daughter cell during mitosis. Two parallel conserved machineries, one dependent on the microtubule associated protein Kar9 and the other on the microtubule motor Dynein, robustly orient the mitotic spindle [1–3]. However, the mitotic spindle might misorient due to environmental conditions or defects in spindle positioning systems. In cells with misoriented spindles, a surveillance mechanism named the spindle position checkpoint (SPOC) halts cell cycle progression in late anaphase of mitosis [4–8]. This way, SPOC provides cells with time to correct the error in spindle positioning and assures correct segregation of genomic DNA before the separation of mother and daughter cells. SPOC impairment causes multiploidy and genomic instability. A SPOC-like surveillance mechanism was shown to exist in fruit fly male germline stem cells. In this system, the checkpoint arrests cells prior to mitosis in response to centrosome misorientation to promote asymmetric cell division [9, 10].

SPOC prevents cell cycle progression by blocking the mitotic exit network (MEN) [11–14]. The MEN is a GTPase-driven signaling cascade that activates Cdc14 - a phosphatase that is essential for mitotic exit through inactivation of mitotic cyclin-dependent kinase (M-Cdk) [15]. Cdc14 is held inactive in the nucleolus, bound to the nucleolar resident protein, Net1. In early anaphase, Cdc14 is transiently released from the nucleolus by the Cdc-fourteen early anaphase release (FEAR) pathway [16]. This pool of partially released Cdc14 plays an important role in spindle formation, rDNA segregation and priming of the MEN but is not sufficient to promote mitotic exit [16–18]. For mitotic exit, Cdc14 must be fully released by the MEN in late anaphase.

The MEN is activated by the small GTPase Tem1, the activity of which is negatively regulated by the GTPase activating protein (GAP) complex composed of Bfa1 and Bub2. MEN components including Bfa1-Bub2, Tem1 and the downstream kinases Cdc15 and Dbf2-Mob1 complex associate with the yeast microtubule-organizing center, namely the spindle pole body (SPB), throughout the cell cycle. The SPB functions as a scaffold that promotes activation of MEN in cells with properly aligned spindles [19, 20].

Upon spindle misalignment, the Bfa1-Bub2 GAP complex must be kept in the active form to promote GTP hydrolysis of Tem1, and hence MEN inactivation [21]. Bfa1-Bub2 GAP activity is controlled through phosphorylation of the Bfa1 subunit. Three kinases were shown to phosphorylate Bfa1: polo like kinase Cdc5, the mother cell associated AMPK family kinase Kin4 and mitotic cyclin dependent kinase M-Cdk1 [22–27]. Cdc5 phosphorylates Bfa1 during anaphase in cells with properly aligned spindles. This phosphorylation reduces the activity of the Bfa1-Bub2 GAP complex *in vitro* and *in vivo*, thereby contributing to MEN activation [22, 25]. Kin4 counteracts the phosphorylation of Bfa1 by Cdc5 and is thus essential for SPOC because it keeps the Bfa1-Bub2 GAP complex in its active form [23, 24]. Recently, M-Cdk was reported to phosphorylate Bfa1 at six residues [26]. A non-phosphorylatable mutant of Bfa1 is SPOC-deficient, implying that M-Cdk keeps the Bfa1-Bub2 GAP complex active. The phosphorylation of Bfa1 by M-Cdk is reverted by the pool of Cdc14 phosphatase activated by the FEAR network [26]. Cdc14 thus acts as an inhibitor of the Bfa1-Bub2 GAP complex and by doing so primes MEN activation after anaphase onset [26]. Kin4, on the other hand, is required to re-activate the Cdc14-inhibited-Bfa1 during anaphase [26].

How the SPOC works on a molecular level is still unclear. Loss of microtubule-cortex interactions promotes Kin4 activity upon Bfa1. However, this is likely not the only factor that contributes to SPOC [26, 28, 29]. The compartmentalization of mitotic exit-activating and -inhibiting factors in daughter and mother cells respectively, was also shown to coordinate mitotic exit with the delivery of a nucleus into the daughter cell body [26, 30]. Yet, how these factors are regulated in a spatial and timely manner remains unclear.

Here, we performed a genome-wide genetic screen to search for novel factors involved in SPOC. We found several proteins and protein complexes in the absence of which SPOC was impaired. These included proteins with a known function in SPOC (Bfa1-Bub2 GAP complex, Kin4, Bmh1 and PP2A protein phosphatase complex) and various factors that had not been implicated in SPOC so far, including the GSK3 family kinase Mck1, the histone deacetylase Hos2 and the SWR1 chromatin remodeling complex (SWR1-C). Among those, we analyzed the role of SWR1-C in SPOC. Our data show that cells lacking SWR1-C were able to initiate the SPOC-response upon spindle mis-orientation; however upon prolonged SPOC arrest, they underwent mitotic slippage. We performed an additional genome-wide genetic screen to identify factors that are essential for mitotic slippage in SWR1-C deficient cells. We show that mitotic slippage requires FEAR, the SAGA complex and proteasome components. We propose that SWR1-C acts through MEN to prevent premature mitotic exit upon checkpoint activation possibly by regulating transcription of multiple factors.

## RESULTS

### A genome-wide genetic screen reveals novel genes critical for SPOC integrity

Overexpression of *KIN4* results in a late anaphase cell cycle arrest that mimics a constitutively active SPOC [23]. Thus, cells overexpressing *KIN4* fail to form visible colonies on agar plates (lethal phenotype). This lethality can be rescued by deletion of SPOC components that act downstream of *KIN4* (Bfa1, Bub2, Bmh1) [23, 24, 31] or those that positively regulate *KIN4* (PP2A-Rts1 and Elm1) [32–34]. In order to find novel SPOC components, we performed a genome-wide genetic screen using the synthetic genetic array (SGA) technology [35]. In this screen, we searched for gene deletions that rescue the lethality of *KIN4* overexpression (See Figure 1A for an outline of the screen). Deletions of the known SPOC components *BFA1*, *BUB2*, *RTS1* and *BMH1* [31–34] were among the top 100 hits of the screen (Figure 1B and Supplementary Table 1).

**Figure 1.**
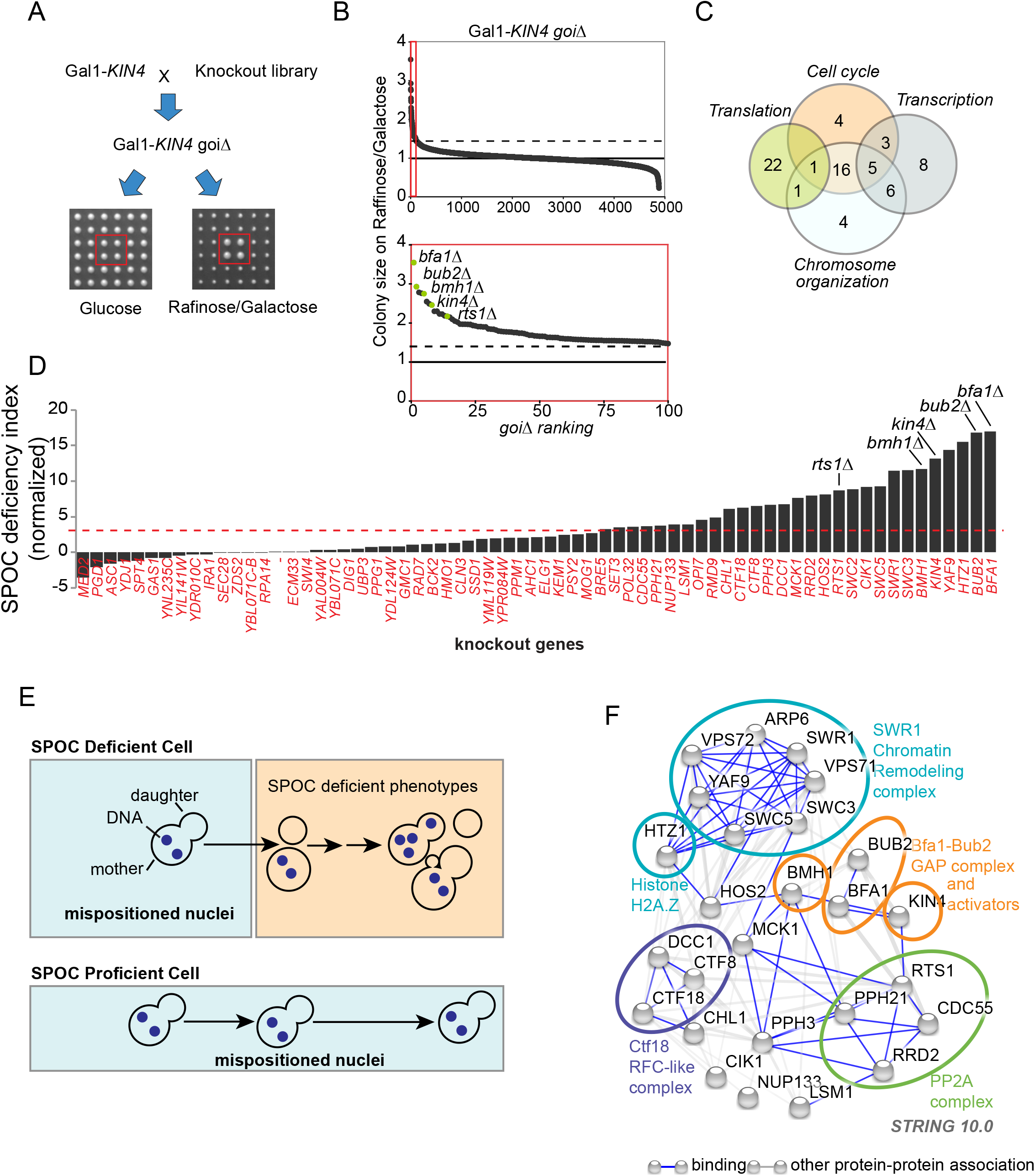
Gene deletions that impair SPOC integrity. **A.** Schematic representation of the screen strategy. The query strain carried a construct for conditional *KIN4* overexpression from the galactose-inducible promoter (Gal1*-KIN4*). The query strain was mated to a sporulated heterozygous diploid knockout library in which each library strain had four technical replicates spotted next to each other (2×2). Obtained diploids were spotted on haploid selection plates and the haploids carrying both the Gal1*-KIN4* and the corresponding gene deletion from the library (*goiΔ*) were selected. These double mutants (Gal1-*KIN4 goiΔ*) were replicated on two types of plates with different carbon sources: glucose (no *KIN4* overexpression) and raffinose/galactose (*KIN4* overexpression). Plates were photographed, and colony sizes were measured in both plates to find colonies that grow under *KIN4* overexpressing conditions. An example of a gene deletion with the desired phenotype is marked with a red box. **B.** Plot showing colony sizes of knockout strains bearing Gal1*-KIN4* on galactose containing agar plates. Knockout strains were ranked from better growing to poorly growing (*goiΔ* ranking from 1 to 4852, where *goiΔ* indicated the deletion of the corresponding gene of interest). The known SPOC proteins among the gene deletions with the top 100 highest colony sizes (red box) are marked in the magnification (lower plot). Dashed line indicates the threshold colony size value of 1.3, while a colony size of 1 is the median colony size on each plate. **C.** Venn-diagram showing the number of genes in indicated GO biological process categories. Remarkably, 70 among the top 100 hits of the screen fell into the indicated categories. **D.** SPOC deficiency indexes of knockouts. Individual gene deletions were combined with the deletion of *KAR9* to induce spindle misalignment. SPOC deficiency was assayed by microscopy of fixed cell populations after DNA staining. 100 cells were counted for knockout per experiment. The graph is a representation of 2 independent experiments. SPOC deficiency index of the *kar9Δ* was normalized to zero. Strains were considered SPOC deficient if the SPOC deficiency index is greater than 3.5 (marked by the red dashed line that is two standard deviations above *kar9Δ* SPOC deficiency). Each bar represents a different gene deletion. Bars corresponding to the known SPOC genes are indicated. Cells with normal aligned, misaligned nuclei as well as the cells with SPOC deficient phenotypes (as depicted in **E**) were counted. SPOC deficiency index represents the ratio of the frequencies of SPOC deficient phenotypes to misaligned cells. **E.** Schematic representation of SPOC deficiency. A SPOC deficient cell with a mispositioned anaphase spindle (indicated by two separated DNA regions in the mother cell compartment) exits mitosis and undergoes cytokinesis giving rise to cells with multiple nuclei and no nucleus. Further divisions of the same cell increase the multiple nuclei phenotype and also cause multi-budded phenotypes (upper panel). SPOC proficient cells, on the other hand, neither exit mitosis nor undergo cytokinesis as long as the spindle stays mispositioned (bottom panel). **F.** Genetic and physical interactions between the proteins found to be involved in SPOC (based on experimental sources in string database). Disconnected nodes were removed. The ovals encircle the proteins that belong to the same protein complex..

The vast majority of the hits fell into Gene Ontology (GO) categories of cell cycle, chromosome organization, transcription and translation (Figure 1C). Several genes related to ribosome biogenesis, tRNA synthesis and galactose metabolism were excluded from further analyses (Figure S1, Supplementary Table 2), reasoning that they may interfere with *KIN4* overexpression from the Gal1 promoter. To validate the remaining hits, we individually deleted each gene in an independent strain background carrying the Gal1*-KIN4* construct. Using growth assays and immunoblotting to measure Kin4 protein levels, we confirmed that deletion of 52 screen hits (genes of interest, *goiΔ*) promoted growth upon *KIN4* overexpression without affecting the extent of overexpression (Supplementary Table 2, Figure S1).

Next, we analyzed SPOC proficiency of *goiΔ* strains (Figure 1D-E, Supplementary Table 2). SPOC proficiency of *goiΔ* strains was assessed in a *kar9Δ* background. Kar9 is a conserved protein involved in spindle positioning [36]. Deletion of *KAR9* causes frequent spindle misalignment that allows assaying of SPOC integrity. The SPOC induces a late anaphase arrest in cells with mis-aligned spindles (i.e. when the mitotic spindle remained in the mother cell body). However, in the absence of SPOC, cells with mis-aligned spindles divide and undergo another round of budding and DNA replication. This leads to accumulation of SPOC-deficient phenotypes such as multi-budded and multi-nucleated cells (Figure 1E). We categorized 26 *goiΔ* strains as SPOC deficient based on the multi-nucleation and multi-budding phenotypes (Figure 1D. These included the known SPOC components and 21 genes that were not yet implicated in SPOC or mitotic exit regulation (Figure 1D). Intriguingly, many of these genes encode subunits of multiprotein complexes (Figure 1F).

### SWR1-C is not required for the metaphase arrest upon microtubule depolymerization

Several subunits of the conserved SWR1 chromatin-remodeling complex SWR1-C were captured in our screen (Figure 1F) [37]. SWR1-C replaces the chromatin bound H2A with the H2A.Z histone variant [38–40]. Interestingly, *HTZ1* (gene that encodes for H2A.Z) was also found in the screen (Figure 1D-F). This suggested that deposition of H2A.Z in the nucleosomes by SWR1-C is necessary for the late anaphase arrest induced by *KIN4* overexpression, as well as the anaphase arrest triggered by the spindle position checkpoint upon spindle misalignment.

We next asked whether SWR1-C also contributes to the mitotic arrest imposed by the spindle assembly checkpoint (SAC). SAC monitors kinetochore-microtubule attachments and prevents anaphase onset upon failure of bipolar kinetochore-microtubule attachment [41]. The SPOC proteins Bfa1 and Bub2 but not Kin4 are required for the mitotic arrest induced by SAC [23, 24, 42–45]. To induce SAC arrest, we treated cells with the microtubule depolymerizing drug nocodazole. Wild type cells were arrested as large budded cells, without degrading Securin/Pds1 (Figure 2). In the absence of *BUB2*, Securin/Pds1 was degraded (time point 90) and cells started re-budding, indicating cell cycle progression (Figure 2). *swr1Δ* cells behaved similar to wild type and *kin4Δ* cells (Figure 2). Therefore, we conclude that SWR1-C is required for SPOC but not for SAC.

**Figure 2.**
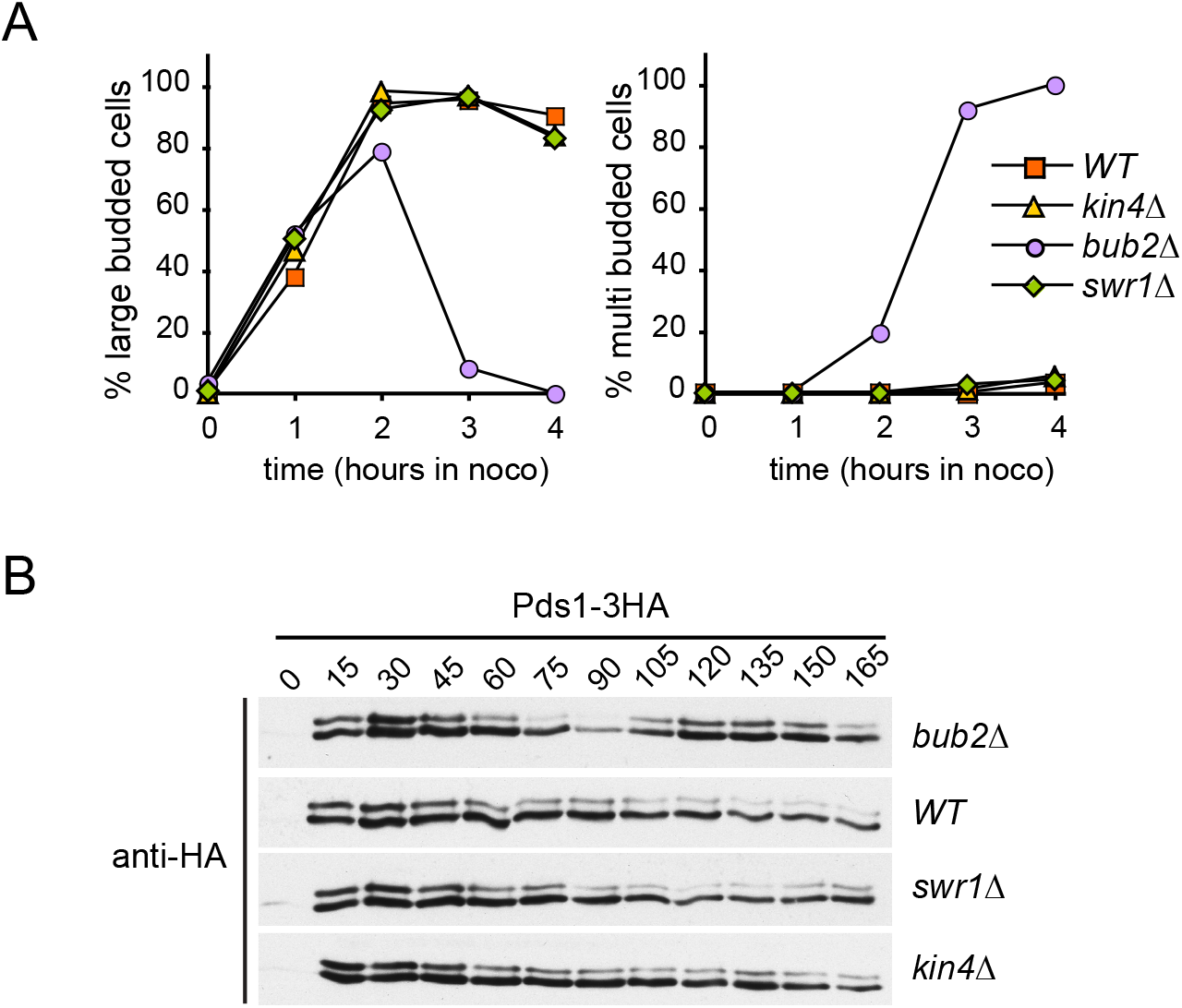
*SWR1-C* is not essential for SAC integrity. **A-B.** SAC integrity of indicated cell types. Cells were released from a G1 block (t=0) into nocodazole containing medium. Samples were collected at indicated time points. (**A**) Percentages of large-budded and multi-budded cells were scored and (**B**) Pds1-3HA levels were analyzed by immunoblotting at the indicated time points (in min.).

### SWR1-C does not influence Kin4 activity or Bfa1 regulation by Kin4

We next investigated whether SWR1-C is required for Kin4-dependent SPOC activation. Hallmarks of SPOC signaling are recruitment of Kin4 to the SPB and phosphorylation of Bfa1 by Kin4 upon spindle misalignment, which subsequently promotes a decrease in the SPB-bound levels of Bfa1 [5, 7, 23, 24, 27, 29, 46, 47]. Deletion of *SWR1* influenced neither Kin4 localization (Figure S2A) nor the ability of Kin4 to phosphorylate Bfa1 in vitro (Figure S2B) or in vivo (Figure S2C). Furthermore, SPB-bound Bfa1 levels decreased in *swr1Δ* cells upon spindle misalignment, similar to control cells (Figure S2D). These data altogether suggest that SWR1-C is not required for Kin4 activity or Kin4-dependent regulation of Bfa1 upon SPOC activation.

### SWR1-C is required for prolonged SPOC arrest

To examine the SPOC in more detail, we compared the duration of anaphase in *swr1Δ kar9*Δ cells upon spindle misalignment and upon correct spindle alignment through time-lapse fluorescence microscopy. *kar9Δ* cells had an anaphase duration of 20 min when their spindle was correctly aligned in the mother to daughter direction, whereas cells with misaligned spindles stayed arrested during the duration of the movie (60 min) with an intact misaligned anaphase spindle (Figure 3A). *kin4Δ kar9Δ* cells with misaligned spindles broke their anaphase spindle with the same timing as cells with normal aligned anaphase spindle [32] (Figure 3A). The vast majority of *swr1Δ kar9Δ* cells broke their spindle despite spindle misalignment (Figure 3A). Nevertheless, anaphase duration was 8 minutes longer in *swr1Δ kar9Δ* cells with spindle misalignment than those with correct spindle alignment. In addition, failure of SPOC in *swr1Δ kar9Δ* cells was not fully penetrant, as 20% of the cells with misaligned spindles were able to stay arrested for a longer time period (Figure 3A). Therefore, unlike *kin4Δ kar9Δ* cells that cannot engage the SPOC, *swr1*Δ *kar9Δ* cells are able to delay mitotic exit upon spindle mis-positioning but the delay is not maintained. Our data thus suggest a role for SWR1-C in preventing mitotic slippage upon prolonged SPOC arrest.

**Figure 3.**
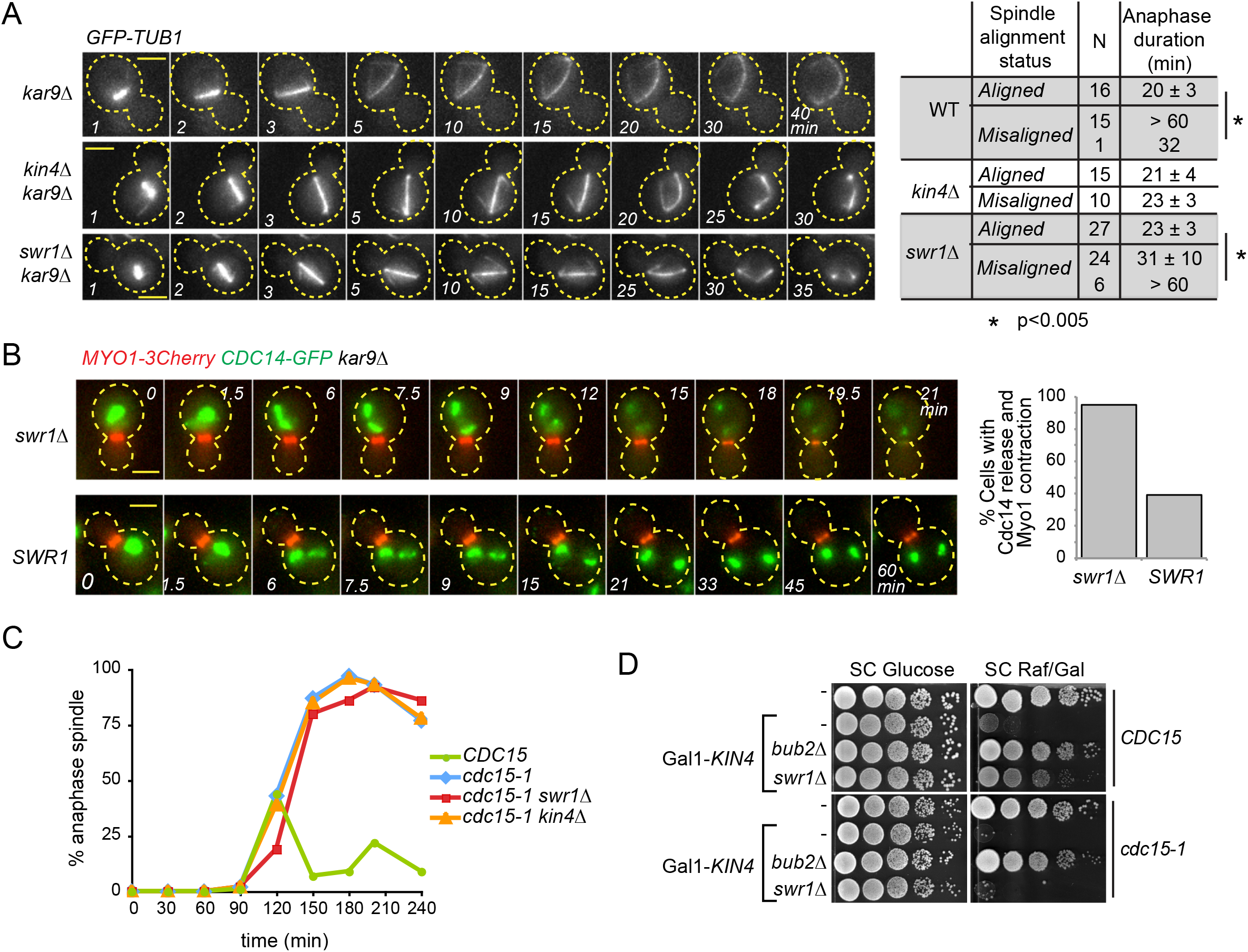
Deletion of *SWR1* causes slippage from the SPOC arrest. **A.** Duration of anaphase in indicated cell types during correct spindle alignment and spindle misalignment. GFP labeled tubulin (*GFP-TUB1*) was monitored by time-lapse microscopy with one-minute time resolution. The duration of anaphase was calculated as the time elapsed from the start of the fast spindle elongation phase (metaphase-anaphase transition) until spindle breakdown. N: number of cells analyzed. Asterisks indicate significant difference according to Student’s t-test (p<0.05). Representative still images of indicated *GFP-TUB1 kar9Δ* cells monitored during spindle misalignment are shown on the left. Cell boundaries are outlined with dashed lines. Arrows indicate the point of spindle breakdown. Time point 1 is one minute before the metaphase-anaphase transition, which coincides with the start of fast spindle elongation phase. Scale bar: 3 μm. **B.** Representative still images from the time-lapse movies of *CDC14-GFP MYO1-3mCherry* bearing *kar9Δ* cells with misaligned spindles. Scale bar: 3 μm. Percentage of cells with misaligned spindles that released Cdc14-GFP and contracted Myo-3mCherry is plotted. N = 20 and 17 for *swr1Δ* and *SWR1,* respectively. **C.** Percentage of cells with intact anaphase spindles. Cultures of indicated cell types bearing GFP-Tubulin were synchronized in G1 by alpha factor treatment (t=0) at the permissive temperature of *cdc15-1* (23 °C) and released in alpha factor free medium at the restrictive temperature (37 °C). Samples were collected at the indicative time points and percentage of cells with intact anaphase spindles (spindle length > 3 μm) was scored. **D.** Serial dilutions of *CDC15* or *cdc15-1* cultures with indicated genotypes were spotted on Gal-*KIN4* repressing (SC Glucose) or inducing (SC Raf/Gal) agar plates.

### Absence of SWR1-C allows MEN activation in cells with misaligned spindles

We sought to understand how the absence of SWR1-C promotes slippage of cells with misaligned spindles out of mitosis. Mitotic exit in budding yeast is triggered by the phosphatase Cdc14 through inhibition of the mitotic cyclin dependent (M-Cdk) kinase activity and dephosphorylation of M-Cdk targets. Employing fluorescence time-lapse microscopy, we asked whether *swr1Δ* cells with misaligned spindles fully activate Cdc14 during spindle misalignment. For this, we scored the release of Cdc14-GFP from the nucleolus in *SWR1 kar9Δ* and *swr1Δ kar9Δ* cells during spindle misalignment. In addition, Myo1-mCherry contraction was analyzed in the same cells as a marker for cytokinesis. The majority of *swr1Δ* cells with misaligned nuclei released Cdc14-GFP from the nucleolus (Figure 3B). Approximately 4 min (4 ± 2 min) after the initiation of Cdc14-GFP release, Myo1-3mCherry started contraction, which lasted ~6 min (6 ± 2min). On the contrary, the majority of the *SWR1* cells with misaligned spindles did not release Cdc14-GFP or contract Myo1-3mCherry. Thus, mitotic slippage imposed by *SWR1* deletion occurs through Cdc14 release.

### SPOC slippage of *swr1Δ* cells requires an active MEN

We then asked whether deletion of *SWR1* could bypass the necessity of MEN for mitotic exit. To address this question, we analyzed the cell cycle progression of a temperature sensitive mutant of the MEN kinase Cdc15 (*cdc15-1*) in the absence and presence of *SWR1*. At restrictive temperature, *cdc15-1* and *cdc15-1 swr1Δ* cells arrested in late anaphase with intact anaphase spindles (Figure 3C). Furthermore, deletion of *SWR1* did not rescue the growth defects of temperature sensitive MEN mutants (Figure S3A). Strikingly, deletion of *SWR1* was not able to rescue the lethality of *KIN4* overexpression in the *cdc15-1* mutant at a semi-permissive temperature (Figure 3D). The same was observed for other MEN temperature sensitive mutants (Figure S3B). These data altogether show that deletion of *SWR1* does not bypass MEN; on the contrary it requires a functional MEN for promoting growth of *KIN4*-overexpressing cells. Taking into consideration that *swr1Δ* cells are able to promote Cdc14 release during spindle misalignment (Figure 3B), we conclude that *SWR1* deletion likely activates a mitotic exit promoting mechanism that acts upstream of MEN.

### Roles of SWR1-C in heterochromatin anti-silencing and DNA damage response do not contribute to inhibition of mitotic exit

Chromatin remodeling by SWR1-C functions in transcriptional regulation through both gene activation and silencing [48, 49]. SWR1-C is also involved in prevention of heterochromatin spreading by antagonizing Sir-dependent silencing [50, 51]. We asked whether Sir-dependent heterochromatin spreading causes the observed growth phenotype of *swr1Δ* cells upon *KIN4* overexpression. Deletion of *SIR2,* which reverses the silencing caused by H2A.Z loss at heterochromatin [50, 51], did not reverse the growth phenotype of Gal1*-KIN4 swr1Δ* (Figure 4A). Thus, *SWR1* deletion rescues the lethality of *KIN4* overexpression through a mechanism different than spreading of silent heterochromatin.

**Figure 4.**
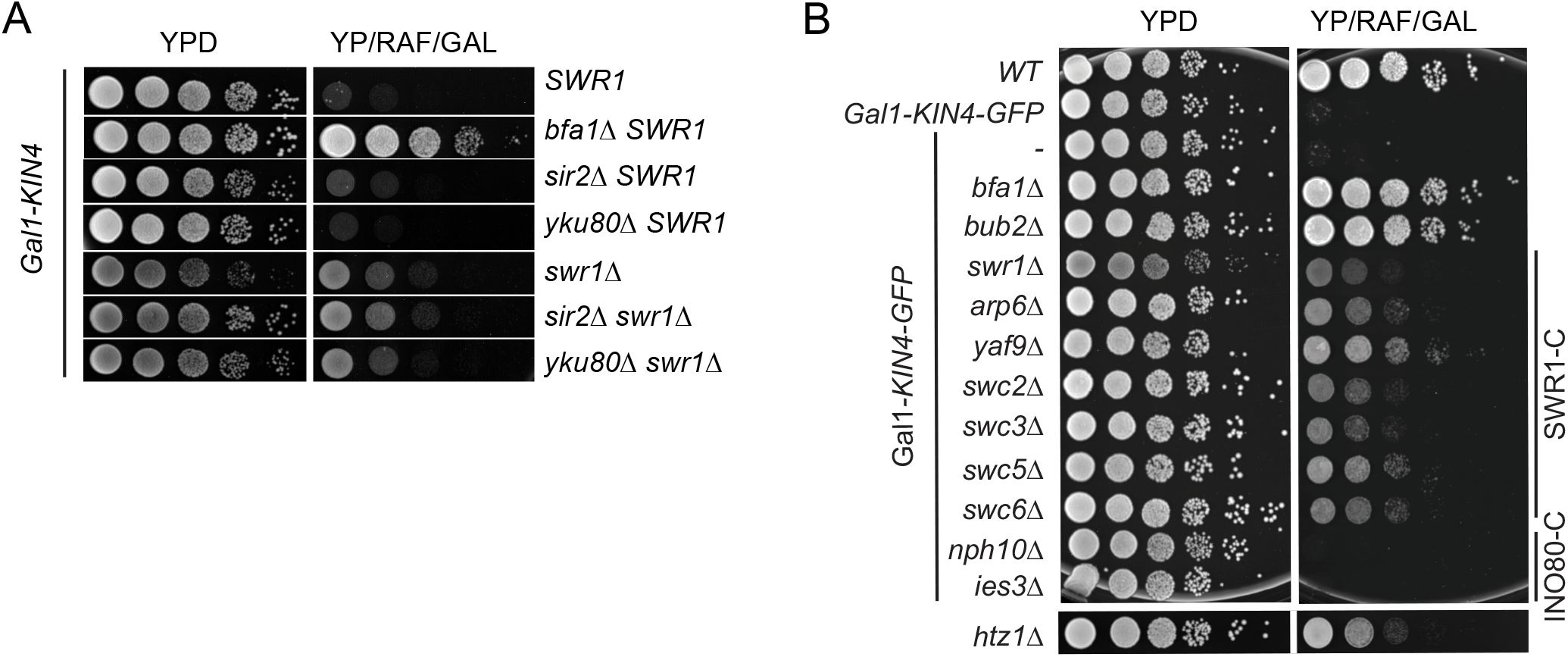
Analysis of the involvement of DNA damage repair, prevention of heterochromatin spreading, perinuclear DNA sequestration or INO80-C related genes in the suppression of *KIN4* overexpression lethality. **A-B.**Serial dilutions of indicated strains were spotted on the indicated agar plates that suppress (YPD) or induce (YP/RAF/GAL) *KIN4* overexpression. (**A**) Function of Swr1 in suppressing Kin4 overproduction does not require Sir2 or Yku80. (**B**) Kin4 overexpression lethality is suppressed by deletions of SWR1-C and *HTZ1* but not INO80-C. *BFA1* or *BUB2* deletion reverts Kin4 overproduction lethality and was used as comparison.

SWR1-C also functions in the DNA damage response through recruitment of Yku80 at double strand breaks to facilitate nonhomologous end-joining (NHEJ) [52]. However, unlike *SWR1*, deletion of *YKU80* did not rescue the toxicity of *KIN4* overexpression (Figure 4A) suggesting that the role of SWR1-C in prevention of mitotic exit is not via its function in NHEJ.

SWR1-C and the INO80 chromatin-remodeling complex (INO80-C) have roles in sequestration of DNA double strand breaks (DSB) to the nuclear periphery [53–55]. INO80-C belongs to the same nucleosome remodeler family as SWR1-C and shares components with SWR1-C [56]. Specific disruption of SWR1-C but not of INO80-C rescued the lethality of *KIN4* overexpression (Figure 4B). Therefore, it is unlikely that the perinuclear DSB sequestration relates to the prevention of mitotic exit in *KIN4* overexpressing cells. Altogether we conclude that the contribution of SWR1-C to the prevention of mitotic exit is not related to its roles in heterochromatin spreading, DNA repair and perinuclear DNA sequestration.

### SPOC slippage requires FEAR, proteasome and SAGA

To determine which gene or gene clusters contributed to the SPOC bypass of *SWR1* deleted cells, we performed a SGA-based genome-wide screen using *swr1Δ* and Gal1-*KIN4 swr1Δ* cells. We reasoned that deletion of genes involved in SPOC bypass would cause death of *swr1Δ* cells overexpressing *KIN4* (Figure S4A). We found 69 gene deletions that caused growth defects specifically in Gal1-*KIN4 swr1Δ* cells on galactose plates (Supplementary Table 3). Most of these genes were in GO categories of transcription, cytoskeletal organization, cell cycle, and cell wall organization (Figure 5A). Among them, many coded for proteins in common protein complexes or proteins participating in common signaling pathways or cellular processes, such as the SAGA complex, PAS complex, mannose transferase complex, proteasome, cell wall integrity pathway, FEAR-pathway, cytokinesis and abscission processes (Figure 5B).

**Figure 5.**
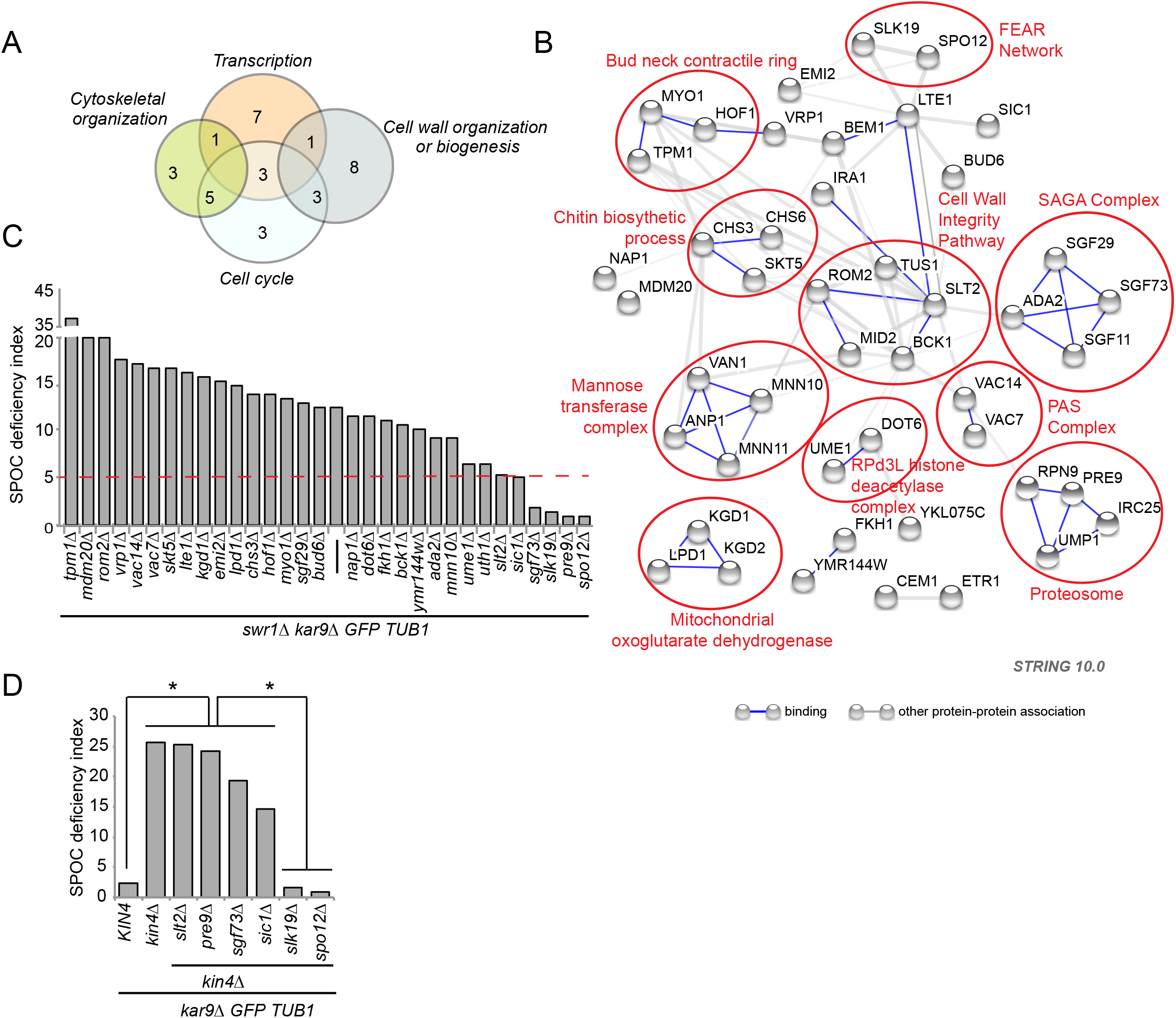
SGA screening identifies genes that are essential for the growth of *swr1Δ* Gal1-*KIN4* cells under *KIN4* overexpressing conditions. **A.** Number of screen hits found within each of the indicated GO categories. **B.** Interaction map of the hits. Circles indicate proteins with related functions. **C-D.** SPOC deficiency indexes of the indicated strains.

In an independent strain background, we performed deletions of at least two genes from each category represented in Figure 5B. We confirmed that 90% of these gene deletions caused growth retardation in *swr1Δ* cells under *KIN4*-overexpressing conditions (Supplementary Table 4, Figure S4B). We next tested whether those gene deletions also affected SPOC in *swr1Δ kar9Δ* cells. Remarkably, SPOC deficiency of *swr1Δ* cells was completely rescued by deletion of four genes. These were the FEAR pathway components *SPO12* and *SLK19,* which enhance MEN activation; the SAGA histone acetyltransferase complex component *SGF73,* which is involved in regulation of RNA Polymerase II transcription preinitiaition and *PRE9*, which is the only non-essential 20S proteasome subunit (Figure 5C). Deletion of the MAPK *SLT2* and M-Cdk inhibitor *SIC1* also greatly retarded formation of multi-nucleated and multi-budded cells, but did not completely rescue the SPOC deficiency of *swr1Δ* cells (Figure 5C). Most of gene deletions that reverted the growth phenotype of Gal1-*KIN4 swr1Δ* cells however did not revert the SPOC deficiency phenotype of *swr1Δ kar9Δ* cells. This implies that the late anaphase arrest in cells with correctly and misaligned spindles might have different characteristics. Interestingly, some gene deletions even caused an increment in accumulation of SPOC deficient phenotypes. Essentially, these genes (*TPM1*, *MDM20, ROM2*) were related to actin organization and cellular polarity [57–60] suggesting that the increased SPOC defect could stem from the inability of spindle re-alignment.

We next asked whether the genes required for SPOC deficiency of *swr1Δ* cells were also required for SPOC deficiency of *kin4Δ* cells. In concordance with previous publications [26, 28], deletion of FEAR network components *SPO12* and *SLK19* rescued SPOC deficiency of *kin4Δ* cells. On the other hand, deletion of *PRE9*, *SLT2, SGF73* and *SIC1* did not significantly impact upon SPOC deficiency of *kin4Δ* cells, albeit a slight reduction due to *SIC1* and *SGF73* deletion. This data suggests that *PRE9*, *SLT2, SGF73* and *SIC1* act by a mechanism different than the FEAR network in SPOC slippage.

Taken together, in addition to the FEAR network, which eventually impinges upon the MEN via Bfa1, Cdc15 and Mob1 regulation, mitotic slippage of SPOC in *swr1Δ* cells require RNA polymerase II-dependent transcription and proteasome-dependent protein degradation. Owing to the role of SWR1-C in gene expression, it is tempting to speculate that SWR1-C dependent transcriptional regulation is important for prevention of mitotic slippage during SPOC arrest.

## DISCUSSION

The spindle position checkpoint (SPOC) exploits proteins localized to the spindle pole bodies (SPBs) as well as mother and daughter cell cortexes to coordinate mitotic exit with the direction of chromosome segregation. How SPOC operates on a molecular level and how it maintains a robust cell cycle arrest remains elusive. In this study, we performed a genome-wide screen that identified novel proteins essential for SPOC integrity. Our analysis uncovered a novel function for the chromatin-remodeling complex, SWR1-C, in preventing mitotic slippage upon prolonged SPOC arrest.

### Identification of novel SPOC components

Higher levels of Kin4 inhibit cell growth due to constitutive inactivation of MEN through the Bfa1-Bub2 GAP complex [23]. Here, we searched for gene deletions that rescued the growth defect of *KIN4* overexpressing cells. Our screen was designed to identify novel genes that inhibited mitotic exit. Among them, we were particularly interested in SPOC-related genes that either directly promoted Kin4 function (e.g. Kin4 regulators or Kin4 substrates) or inhibited mitotic exit downstream or in parallel to *KIN4* (e.g. inhibitors of MEN). To find the SPOC regulators among those putative mitotic exit inhibitors, we performed a secondary screen using cells that lacked the spindle-positioning protein, Kar9. This screen revealed that the majority of genes identified as potential mitotic exit inhibitors were indeed important for SPOC functionality. The known SPOC regulators as well as novel SPOC components were among the hits of the secondary screen. These novel SPOC components included a protein kinase (Mck1), a protein phosphatase (Pph3) and chromatin related proteins such as a histone deacetylase (Hos2), and a chromatin remodeler (SWR1) among others. Importantly, SWR1 chromatin remodeling complex (SWR1-C) components constituted the majority of novel SPOC proteins found in the screen. SWR1-C remodels the chromatin by replacing histone H2A with the histone variant H2A.Z at the chromatin bound nucleosomes. The *HTZ1* gene, which encodes for the H2A.Z histone variant, was also identified in our screen as a novel SPOC component. In this study, we investigate the function played by SWR1-C in SPOC in more detail. The function of other novel SPOC proteins will be in the scope of other studies.

### SWR1-C prevents mitotic slippage during SPOC arrest

Cells lacking SWR1-C components or the histone H2A.Z accumulated multiple nuclei when spindle alignment was impaired. Time-lapse analysis of *swr1Δ* cells further established that the majority of SWR1-C deficient cells with misaligned spindles exited mitosis, which indicates the failure of SPOC in the absence of SWR1-C function. A closer look at anaphase duration in cells lacking SWR1-C showed that anaphase lasted eight minutes longer in cells with misaligned spindles compared to the cells with correct spindle alignment. Thus, SWR1-C deficient cells can indeed delay mitotic exit for about eight minutes in response to spindle misalignment, however they cannot maintain a longer arrest.

The immediate response to SPOC activation is the recruitment of Kin4 kinase to both SPBs, the phosphorylation of Bfa1 by Kin4 and the change in Bfa1 localization from asymmetric (strong at one SPB) to symmetric (weak at both SPBs). These events are essential for Tem1 inactivation that is indispensable for a late anaphase arrest [61]. Analysis of the SPOC mechanism verified that SWR1-C deficient cells were in fact able to initiate these SPOC-responses upon spindle mis-orientation. Yet, they later released Cdc14 and constricted their actomyosin ring, indicating that the cells were able to exit mitosis and undergo cytokinesis despite Bfa1-Bub2 activation. Thus, inactivation of Tem1 by Bfa1-Bub2 is essential but not sufficient to hold mitotic exit for a prolonged time. Our data indicate that a robust SPOC arrest requires SWR1-C. Altogether these data led us propose two levels of SPOC response; first an “immediate response” that results in inactivation of Tem1 by the core SPOC mechanism involving Bfa1-Bub2 and Kin4 and second a “late response” that prolongs the arrest dependent on SWR1-C. Such dual level regulation might resemble the rapid and slow responses to DNA damage. In case of the DNA damage checkpoint, a rapid response involves degradation of the CDK activating phosphatase Cdc25, whereas the slow response involves transcriptional activation of targets including the CDK inhibitor (CKI) p21 [62].

Genetic analysis indicated that functions of SWR1-C in telomere maintenance or DNA damage response are not relevant to our observations. In addition, our data showed that mitotic slippage that takes place in the absence of SWR1-C is dependent on SAGA complex, which is another histone modifying protein complex bearing histone acetyl transferase and deubiquitination functions that are important for transcriptional activation, particularly through RNA PolII dependent elongation [63]. Thus, SWR1-C dependent regulation of gene expression might be required for the late SPOC response.

Chromatin is highly compacted during mitosis and most gene regulatory elements are cleared from this condensed chromosome. Therefore, it has been generally thought that transcription is completely silenced during mitosis and activated during mitotic exit. However recent studies have proven that RNA PolII remains active on the chromatin and a low level transcription persists during mitosis in a promoter dependent manner [64–67]. Thus, it is plausible that transcriptional regulation may as well regulate mitotic progression in late mitosis. Indeed, microarray analysis of SWR1-C dependent gene expression in a late anaphase arrest (*cdc15-as*) showed that transcriptional profiles of late anaphase-arrested cells differed in the presence and absence of SWR1-C (our unpublished data). We reasoned that SWR1-C might regulate transcription in anaphase-arrested cells. However, using this data we failed to identify individual genes or group of genes involved in the late anaphase arrest of cells with or without spindle misalignment. More needs to be done to understand the contribution of transcriptional regulation on control of mitotic exit.

### SWR1-C does not inhibit FEAR or MEN directly

FEAR is a prominent pathway that impinges on MEN activity via partial release of Cdc14 from the nucleolus during early anaphase. Previous studies showed that FEAR activation promotes mitotic exit and contributes to mitotic slippage of cells with misoriented spindles [30]. We initially thought that SWR1-C might directly regulate Cdc14 release from the nucleolus. However, this was not the case. Cdc14 partial release did not take place prematurely in *swr1Δ* cells (our unpublished observation). Furthermore, mitotic slippage of *swr1Δ* cells that overexpressed *KIN4* required the presence of a functional MEN and FEAR pathways. Of importance, *SWR1* was required for the SPOC but not for the SAC-mediated mitotic arrest. Therefore mitotic slippage that occurs in the absence of SWR1-C seems to be restricted to the anaphase of mitosis. These data altogether are consistent with our model that SWR1-C maintains the late anaphase arrest after the immediate SPOC response that inhibits MEN at the level of Tem1 in cells with misaligned spindles.

### Factors required for mitotic slippage in the absence of SWR1-C

Using an unbiased genetic approach, we identified non-essential genes that contributed to mitotic slippage of SWR1-C deficient cells with misaligned spindles. Most of these genes were factors already known to be involved in mitotic exit. For example, *SIC1*, the mitotic CKI, is important for M/G1 transition [68]. Likewise, the proteasome (Pre9 being one of the non-essential component of 20S proteasome) is required for degradation of the mitotic cyclin Clb2 to achieve mitotic CDK inactivation [69]. The FEAR pathway (*SPO12* and *SLK19* being FEAR components), on the other hand, is crucial for priming mitotic exit at several levels, such as at the level of Bfa1-Bub2 inactivation and, Cdc15 and Dbf2/Mob1 localization at SPBs [17, 18, 26]. Importantly, deletion of *SPO12* or *SLK19* reverted the SPOC deficiency of *swr1Δ* cells as well as *kin4Δ* cells [26, 28], highlighting the role of FEAR pathway in bypassing the immediate SPOC response. Unlike the FEAR pathway, the impairment of the proteasome and deletion of the mitotic CKI reverted the SPOC deficiency in *swr1Δ* but not *kin4Δ* cells. Thus, mitotic slippage that occurs in the absence of SWR1-C particularly requires mitotic CKI and proteasome. Taken together, the late SPOC response mediated by SWR1-C might involve a regulation at the level of the Sic1 CKI or the proteasome. Puzzling, we could not detect any difference in Sic1 or Clb2 mRNA levels during a late anaphase arrest with or without SWR1-C (data not shown). Therefore, if SWR1-C directly regulates *SIC1* or *CLB2* it is likely not at the level of *SIC1* or *CLB2* transcription.

In addition, we found that components of the cell wall integrity (CWI) pathway contributed to the SPOC deficiency of *swr1Δ* cells. CWI pathway regulates G1/S transition and DNA replication in response to cell wall damage [70, 71]. CWI pathway was also shown to stabilize Sic1[71]. However, we could not detect activation of the CWI pathway in Gal1-*KIN4* or Gal1-*KIN4 swr1Δ* cells upon induction of *KIN4* overexpression (data not shown). Therefore, how CWI might contribute to mitotic exit of *swr1Δ* cells remains unclear.

We furthermore found components that belong to the histone acetyl transferase and histone deubiquitylating modules of the SAGA complex to be crucial for mitotic exit in the absence of SWR1-C. Among those, *SGF73* a component of the histone deubiquitylating module (DUBm) was also necessary for mitotic slippage of *swr1Δ* cells with a mispositioned spindle. DUBm of SAGA is required for gene activation through RNA PolII [72]. Thus it is tempting to speculate that genes activated by SAGA in the absence of SWR1-C might be the reason for mitotic slippage.

### Chromatin remodelers and checkpoints

Chromatin remodelers implicated in checkpoint control is not limited to SWR1-C. For example, Isw2 and Ino80 chromatin remodelers facilitate S phase checkpoint deactivation by directly interacting with Replication Protein A [73]. SWI/SNF chromatin remodeling complex regulates DNA damage checkpoint by activating the checkpoint kinase Mec1 (ATR) through interaction of the Snf2 ATPase subunit with Mec1 [74]. In addition, Ies4 subunit of Ino80 chromatin remodeling complex is a downstream target of the Mec1/Tel1 (ATM/ATR) kinases [75]. Several other ATP dependent chromatin remodelers are also involved in DNA damage response [76]. Considering that DNA replication/damage checkpoints detect abnormalities on the DNA, it is not surprising that the chromatin remodelers participate in this process. On the other hand, data on the function of chromatin remodelers in mitotic checkpoints has so far been limited to the RSC chromatin remodeling complex associated with its Rsc2 subunit which contributes to mitotic slippage of cells from the metaphase arrest triggered by the SAC [77]. In this case, Rsc2 physically interacts with the polo like kinase Cdc5 and affects Net1 phosphorylation to promote Cdc14 partial release from the nucleolus during early anaphase. Thus, RSC chromatin remodeling complex promotes mitotic slippage of metaphase-arrested cells. Importantly, Rsc2 interacts with Cdc5 and hence it is likely that RSC regulates mitotic exit independently of its role in transcriptional regulation. Our finding that SWR1-C plays a critical role in preventing mitotic slippage of SPOC-arrested cells reinforces the importance of chromatin remodelers in mitotic control and opens up new insights into mitotic checkpoint regulation.

## MATERIALS AND METHODS

### Yeast methods, strains and growth

Yeast strains used in this study are isogenic with S288C. The strains constructed for validation of the SGA screen are indicated in Supplementary Tables 2 and 4. Other strains are listed in Supplementary Table 5. PCR-based methods were used for gene deletions and epitope tagging [78, 79]. Genes of interest were expressed from their endogenous promoter unless otherwise stated. Basic yeast methods and growth media were as described [80]. For induction of the Gal1 promoter, yeast extract peptone medium containing raffinose (3%) and galactose (2%) (YP-Raf/Gal) was used. To synchronize the cells in G1-phase, 10 μg/ml synthetic alpha-factor (Sigma, St. Louis, MO) were added to logarithmically growing (log-phase) cultures. For nocodazole treatment (metaphase arrest), 15 μg/ml nocodazole (Sigma) was added to the culture media. Genetic interactions with temperature sensitive mutants and Gal1-*KIN4* strains were evaluated at restrictive temperature and under Gal1 overexpressing conditions, respectively. Growth was assayed by performing drop tests in which serial dilutions of cultures were spotted on corresponding agar plates. Plates were incubated at appropriate temperature for 2-3 days.

### Genome-wide genetic screening

Screens were performed as previously described [81] using heterozygous diploid yeast deletion collection [82] where each strain carries a deletion of a single non-essential gene. Library was spotted on plates in 1536-colony format using a ROTOR colony pinning robot (Singer Instruments, Somerset, UK). Each library strain had four technical replicates spotted next to each other (2×2). Query strains were; AKY1296 (Y8205 Gal1*-KIN4*), AKY1307 (Y8205 Gal1-*KIN4 swr1Δ*) and AKY1346 (Y8205 *swr1Δ*). The deletion collection was sporulated and mated with each of the query strain. Then, colonies were sporulated and haploids carrying simultaneously the query mutations and a gene deletion from the deletion collection (*goiΔ,* goi=gene of interest) were selected. All steps until here were carried on glucose containing plates. Then, plates were replicated either on raffinose/galactose or glucose containing agar media. Glucose containing plates were photographed after one day of incubation at 30°C, whereas Raffinose/Galactose containing plates were photographed after one, two, three and four days of incubation at 30°C (d1, d2, d3, d4 in Table S1). Colony sizes, normalized to the median of each plate, were determined from the photographs using SGATools [83]. For each spot, mean and median colony sizes were calculated from the four technical replicates.

The cross with Gal1*-KIN4* carrying strain was used to find out gene deletions that rescued *KIN4* overexpression toxicity. Hits were selected employing a criteria that included colonies with “median colony size > 1.3” and “standard error of the mean < 25%” on Raffinose/Galactose plates (day 2); and excluded the colonies with “median colony size < 0.3” and “colony counts < 4” on Glucose plates. Among such colonies, those with the top 100 highest median colony size were considered as hits.

Crosses with Gal1-*KIN4 swr1Δ* and *swr1Δ* were used to find out gene deletions that revert the growth rescue of *KIN4* overexpression lethality by *SWR1* deletion. To select the hits, we calculated the ratio between the median colony sizes of *swr1Δ goiΔ* and Gal1-*KIN4 swr1Δ goiΔ* on Raffinose/Galactose plates (day 3). Hits were considered as positives when “median colony size of *swr1Δ goiΔ* / Gal1-*KIN4 swr1Δ goiΔ* > 0.16” and “standard deviations < 0.2” and “colony counts > 2”.

### Analysis of checkpoint integrity

For measurement of SPOC integrity, log-phase *kar9*Δ cells cultured at 23°C were incubated at 30°C for 3-5 h. Cells were fixed using 70% Ethanol and re-suspended in PBS containing 1 μg/ml 4’,6-diamino-2-phenylindole (DAPI, Sigma) and analyzed by microscopy. *kar9*Δ cells bearing *GFP-TUB1* were fixed in 4% paraformaldehyde for 10 min at room temperature. Cells with normal and misaligned nuclei, and cells with SPOC-deficient phenotypes (multiple nuclei in one cell body or single nuclei in a multi-budded cell) were counted based on images were taken in phase contrast and corresponding fluorescence channels. At least 100 cells were count per strain per experiment or time point. Analysis of each strain was repeated in 3 independent experiments. SPOC deficiency index was calculated using the following formula; SPOC deficiency index = (% SPOC deficient phenotypes) / (% misaligned spindle) X 10.

SPOC deficiency index (normalized) = SPOC deficiency index of the *kar9*Δ strain bearing the indicated gene deletion – SPOC deficiency index of *kar9*Δ strain. The functionality of the spindle assembly checkpoint was analyzed upon microtubule depolymerization by nocodazole treatment. Briefly, cells synchronized in G1 using alpha factor were released in nocodazole containing media. Samples were collected for microscopy and protein extract preparation. Samples for microscopy were fixed in 70% ethanol and nuclei were stained with DAPI. Number of nuclei per cell and budding status of at least 100 cells were recorded. Only the percentages of large budded and multi-budded cells were plotted. The degradation of Pds1 was analyzed by immunobloting.

### Fluorescence Microscopy

For time-lapse experiment,s cells were adhered on glass-bottom dishes (MatTek, Ashland, MA) using 6% concanavalin A-Type IV (Sigma). Live-cell imaging were performed using a DeltaVision RT wide-field fluorescence imaging system (Applied Precision, Issaquah, WA) equipped with a quantifiable laser module, an OlympusIX71 microscope with plan-Apo×100NA 1.4 oil immersion objective (Olympus, Tokyo, Japan), a camera (Photometrics CoolSnap HQ; Roper Scientific, Tucson, AZ) and SoftWoRx software (Applied Precision) as previously described [29, 31]. Still images of live or fixed cells were acquired using a Zeiss Axiophot microscope equipped with a 100x NA 1.45 Plan-Fluor oil immersion objective (Zeiss, Jena, Germany), Cascade 1K CCD camera (Photometrics, Tucson, AZ) and MetaMorph software (Universal Imaging Corp., Chesterfield, PA). All images were processed in ImageJ (NIH, Bethesda, MD), Adobe Photoshop CS3 and Adobe Illustrator CS3 (Adobe Systems, San Jose, CA). No manipulations were performed other than brightness, contrast and color balance adjustments.

### Calculation of anaphase duration

Anaphase duration of cells with correct and misaligned spindles was determined from time-lapse series of *GFP-TUB1 kar9Δ* cells. Cells grown at 23°C were analyzed by live-cell imaging at 30°C for 1-1.5 h with 1 min time intervals. The time from the start of fast spindle elongation (metaphase-to-anaphase transition) until spindle breakdown was calculated as anaphase duration [84].

### Protein methods

Yeast protein extracts and immunoblotting were performed as described [78]. Antibodies were mouse anti-Tubulin (TAT1, Sigma), mouse anti-HA (12CA5, Sigma), rabbit anti-Clb2 and guinea pig anti-Sic1 [42]. Secondary antibodies were goat anti-mouse, goat anti-rabbit and goat anti-guinea pig IgGs coupled to horseradish peroxidase (Jackson ImmunoResearch Laboratories Inc, West Grove, PA). In vitro kinase assays of immunoprecipitated Kin4-6HA was performed using MBP-Bfa1 purified from *E. coli* as described previously [42, 85, 86].

## ACKNOWLEDGEMENTS

We would like to thank Astrid Hofmann and Dorothee Albrecht for excellent technical assistance, Elmar Schiebel for strains and access to microscopes. This work was funded by the Deutsche Forschungsgemeinschaft (DFG) Collaborative Research Center SFB1036 (Project TP21 to G.P and Project TP10 to M.K. and A.K.) and DFG Project grant PE1883/1 granted to G.P.; A.K.C. was supported by the German Research Council (DFG, PE1883/2), MSCA Individual Fellowship (796599, COHEMEX) and EMBO-IG (3918). The work of G.P. is supported by the Heisenberg program (PE1883/3) of the DFG.

